# Differences in 5’untranslated regions highlight the importance of translational regulation of dosage sensitive genes

**DOI:** 10.1101/2023.05.15.540809

**Authors:** Nechama Wieder, Elston N. D’Souza, Alexandra C. Martin-Geary, Frederik H. Lassen, Jonathan Talbot-Martin, Maria Fernandes, Sonia P. Chothani, Owen J.L. Rackham, Sebastian Schafer, Julie L. Aspden, Daniel G. MacArthur, Robert W. Davies, Nicola Whiffin

## Abstract

**Background:** Untranslated regions (UTRs) are important mediators of post-transcriptional regulation. The length of UTRs and the composition of regulatory elements within them are known to vary substantially across genes, but little is known about the reasons for this variation in humans. Here, we set out to determine whether this variation, specifically in 5’UTRs, correlates with gene dosage sensitivity.

**Results:** We investigated 5’UTR length, the number of alternative transcription start sites, the potential for alternative splicing, the number and type of upstream open reading frames (uORFs) and the propensity of 5’UTRs to form secondary structures. We explored how these elements vary by gene tolerance to loss-of-function (LoF; using the LOEUF metric), and in genes where changes in dosage are known to cause disease. We show that LOEUF correlates with 5’UTR length and complexity. Genes that are most intolerant to LoF have longer 5’UTRs (*P*<1×10^−15^), greater TSS diversity (*P*<1× 10^−15^), and more upstream regulatory elements than their LoF tolerant counterparts. We show that these differences are evident in disease gene-sets, but not in recessive developmental disorder genes where LoF of a single allele is tolerated.

**Conclusions:** Our results confirm the importance of post-transcriptional regulation through 5’UTRs in tight regulation of mRNA and protein levels, particularly for genes where changes in dosage are deleterious and lead to disease. Finally, to support gene-based investigation we release a web-based browser tool, VuTR (https://vutr.rarediseasegenomics.org/), that supports exploration of the composition of individual 5’UTRs and the impact of genetic variation within them.

## Background

Untranslated regions (UTRs) are the regions flanking the protein-coding sequence of genes that form part of the mRNA, but are not translated into protein. UTRs are important mediators of post-transcriptional regulation, controlling mRNA stability, cellular localisation and the rate of protein synthesis [1]. UTRs are known to vary substantially across genes, both in size, and in the composition of regulatory elements within them. These elements can be linear or structural and often mediate their effects through binding to various proteins and non-coding RNAs [2].

The length of 5’UTRs varies between genes and they can be over 2000 base pairs (bp) long [1]. 5’UTRs of genes where heterozygous loss-of-function (LoF) variants cause developmental disorders (DD) are longer and have more introns than all genes [3]. Alternative splicing within the UTRs occurs in transcripts of at least 13% of mammalian genes [4,5], which may exert another level of post-transcriptional control.

Upstream AUG (uAUG) codons are commonly observed within 5’UTRs [1]. uAUGs can be recognised by the scanning 43S ribosomal subunit and its associated initiation factors leading to the initiation of translation. The prospect of a uAUG initiating translation is dependent on several features such as local sequence context (with a stronger match to the Kozak consensus associated with higher levels of translation [6,7]), position of uAUG within the 5’UTR, and presence of nearby secondary structures in the mRNA [8]. These features influence whether the 43S scans past a uAUG or initiates translation from it. uAUGs are conserved to a significantly greater degree than any other triplet in 5’UTRs [9] and there are fewer uAUGs present in the human genome than would be expected by chance [10].

Translation from a uAUG may have one of multiple effects (Figure 1A). Upstream open reading frames (uORFs) are encoded when an uAUG has an in-frame stop codon within the 5’UTR. If there is no in-frame stop codon, an oORF (overlapping ORF) is formed, whose corresponding stop codon extends beyond the coding sequence (CDS) start. oORFs can either be in-frame with the CDS, resulting in an elongated transcript (N-terminal extension, NTE), or out-of-frame, terminating within the CDS [10,12,13]. Start-stops are uAUGs that are immediately followed by a stop codon, with no codons in between. Start-stops are thought to cause ribosome pausing without the energy-expensive peptide production of uORFs [11,14,15]. It is estimated that half of all proteincoding genes contain at least one uORF, and that active translation of a uORF can reduce downstream translation by up to 80% [13]. Genes with uORFs have been demonstrated to have lower protein expression levels than genes without uORFs in multiple human tissues [16], but recent studies have shown that uORF translation is generally positively regulated with translation of the CDS [17–19].

**Figure 1:**
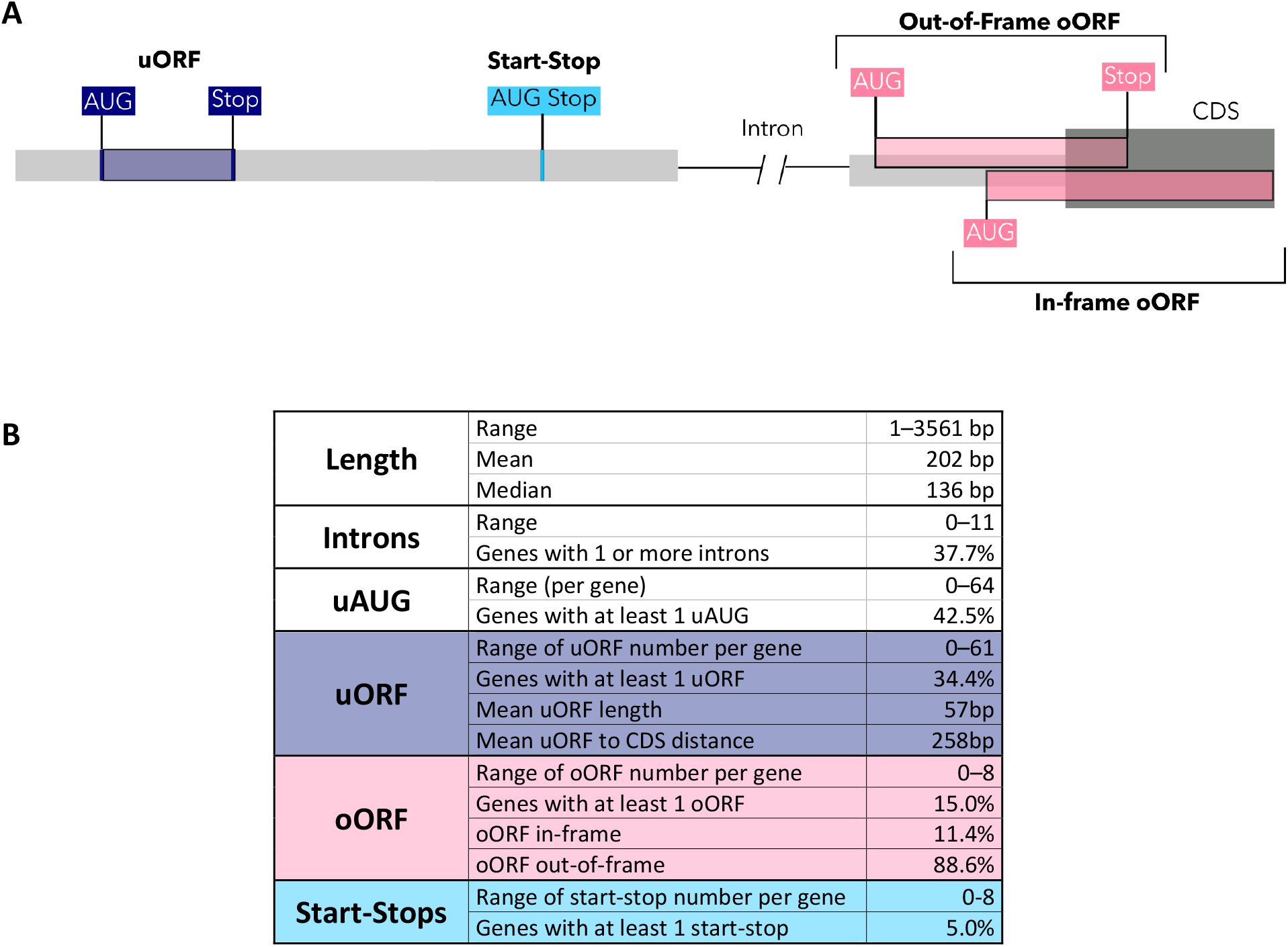
An overview of 5’UTR structure and features. **A)** Illustration of the different types of uAUGs. **B)** Descriptive statistics of annotated 5’UTRs across 18,764 MANE Select transcripts. The length, number of introns, and number and types of uAUGs range widely across genes. uORFs: upstream open reading frame; oORF: overlapping open reading frame; uAUG: upstream AUG (start codon).

Ribosome profiling (Ribo-seq) is an experimental method for determining actively translated regions of the transcriptome, including uORFs [20,21]. Ribo-seq has shown that near-cognate codons (i.e those that differ from AUG by only a single base, such as CUG and GUG) can also act as functional uORF initiation sites [9,12]. After translating a uORF, the ribosome may reassemble and translate the CDS [22]. The efficiency of this ribosome reinitiation has been observed to decrease as the distance between the end of a uORF and start of the CDS decreases [22,23], as the scanning ribosomal subunit requires time (and therefore space) to reacquire a charged tRNA_i_^Met^ with which to recognise the next start codon [9]. uORFs have been found to be depleted in the 100 bp region immediately upstream of the CDS, suggesting that uORFs close to the CDS are selected against as they are more repressive [23].

Genes differ in their tolerance to increases and or decreases in expression levels, or dosage sensitivity. The Genome Aggregation Database (gnomAD) has classified protein-coding genes along a continuous spectrum that represents tolerance to inactivation, termed the “loss-of-function observed/expected upper bound fraction” (LOEUF) score [24]. Our previous work has shown that variants that create uAUGs or disrupt uORFs are under stronger negative selection in genes that are intolerant to loss-of-function [25]. Furthermore, these variants have been shown to cause haploinsufficient disease [3].

Whilst 5’UTRs are known to vary widely in length and composition between different genes, these differences have not been systematically assessed in genes with differing tolerance to changes in dosage. A better understanding of the make-up of 5’UTRs, and the genes for which translational regulation is most critical, is essential to interpreting the impact of genetic variation within these important regulatory elements. Here we systematically analyse 5’UTR regulatory features across and between deciles of LOEUF and in disease gene sets. Our results show that genes which are intolerant to LoF have more complex 5’UTRs that are enriched for cis-acting regulatory elements (including uAUGs). This demonstrates the important role of 5’UTRs in tight regulation of protein levels, particularly for genes where changes in dosage are deleterious and lead to disease.

## Results

### 5’UTRs vary widely across human genes

We analysed 18,764 5’UTRs annotated by the MANE project (v1 MANE Select transcripts) [26]. Of note, 298 (1.6%) MANE Select transcripts do not have an annotated 5’UTR and were excluded. We calculated the overall length of each 5’UTR as well as the position of uAUGs, and introns. The length of 5’ UTRs varies widely between genes, ranging from 1-3,561bp. The number of uAUGs ranges from 0-64 per gene, with 42.5% of 5’UTRs having at least one uAUG (Figure 1B). We further classified these uAUGs by effect, finding that 34.4%, 15.0%, and 5.0% of 5’UTRs contain at least one uORF, oORF, and start-stop element, respectively (Figure 1B).

In addition to annotating ‘predicted uORFs’ as all occurrences of canonical AUG triplets with an in-frame stop codon in each 5’UTR, we used a set of 5,052 functionally validated uORFs detected through ribosome profiling of six cell types and five tissues (‘Ribo-Seq uORFs’) [17]. 1,430 (28.3%) of the predicted uORFs are detected as translated in the Ribo-Seq uORFs dataset (Supplementary Figure 1). In addition, the Ribo-Seq uORF set contains 2,288 additional uORFs that start at non-canonical (non-AUG) start-codons (45.3% of the Ribo-Seq uORFs). Overall, 20.9% of 5’UTRs contain one or more Ribo-Seq uORFs (range 1-11).

### Genes intolerant to loss of function have longer and more complex 5’UTRs

To investigate how 5’UTRs vary by gene sensitivity to decreases in dosage we used LOEUF scores to bin genes into deciles of intolerance to LoF. The lowest deciles represent the genes most intolerant to LoF and the higher deciles represent those most tolerant [24]. We assessed 5’UTR features across LOEUF deciles. For statistical tests, we compared the lowest and highest LOEUF quintiles.

5’UTR length increases with decreased tolerance to LoF (Figure 2A), with the 5’UTRs of genes in the lowest LOEUF quintile being significantly longer than those in the highest LOEUF quintile (mean length 269 bp vs 162 bp; Wilcoxon *P*<1×10^−15^). In other words, genes that are intolerant to LoF have significantly longer 5’UTRs. Given that LOEUF is correlated with CDS length, with shorter genes having less confident LOEUF estimates, we repeated this analysis after removing genes within the bottom 10% of CDS length. Our results remained significant (Supplementary Figure 2; Wilcoxon *P*<1×10^−15^).

**Figure 2:**
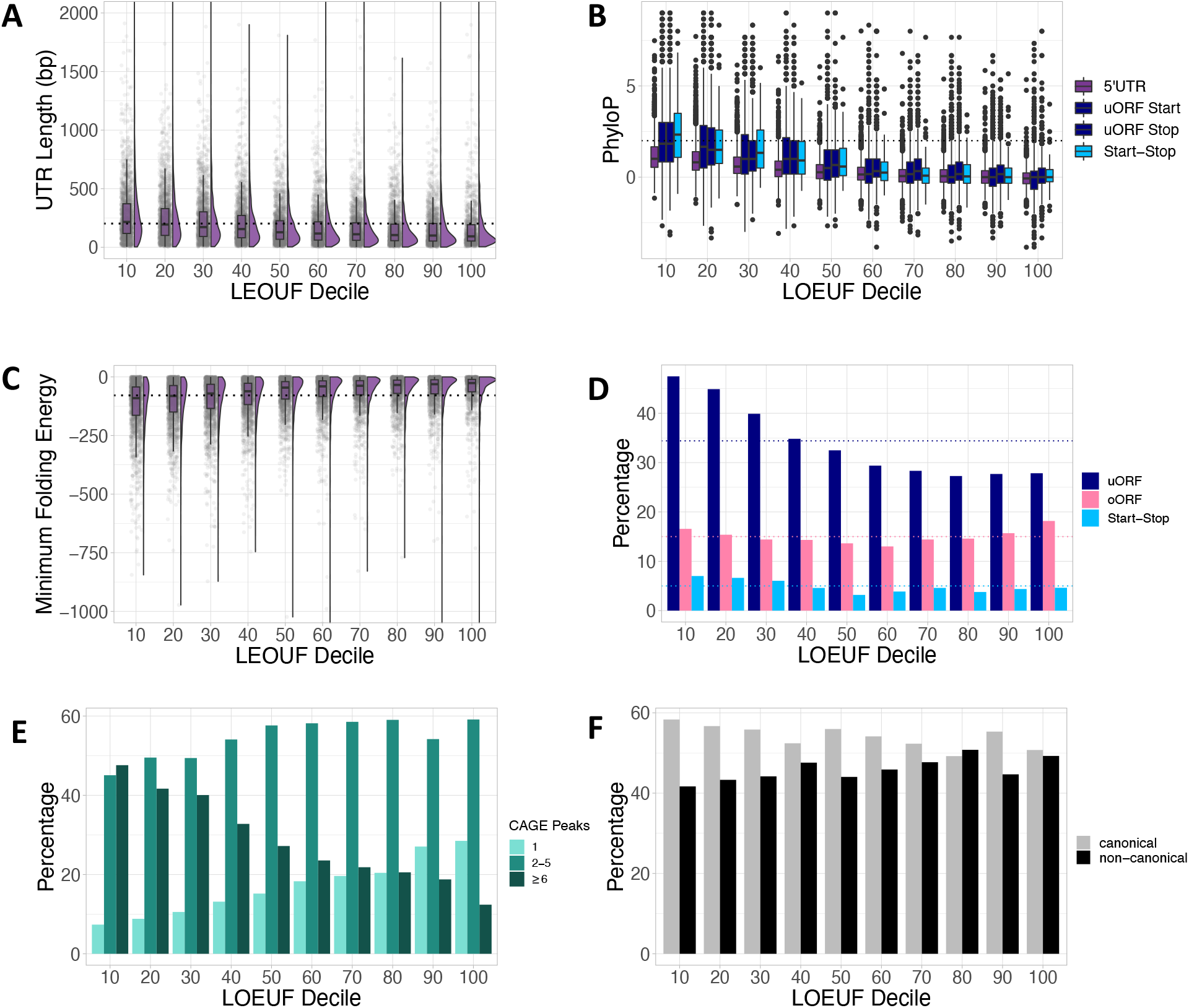
Genes intolerant to LoF have longer and more complex 5’UTRs. **A)** 5’UTRs increase in length with decreasing tolerance to LoF (Wilcoxon P<1×10^−15^). The average 5’UTR length across all genes (202 bp) is shown by a dotted line. The y-axis was truncated at 1,500 bp (39 genes had 5’UTRs >1,500 bp). **B)** The 5’UTRs of genes most intolerant to LoF are more conserved. Average PhyloP scores are plotted for 5’UTRs, uORF start codons, uORF stop codons and start-stops. The dotted line denotes PhyloP=2. **C)** The 5’UTRs of genes most intolerant to LoF have lower minimum free energy (MFE) scores, representing a higher propensity to fold and create structured mRNAs (Wilcoxon P<1×10^−15^). The average MFE across all 5’UTRs is shown as a dotted line (−78.8). The y-axis was truncated at -1,000 (6 genes had MFE <-1000). **D)** Genes most intolerant to LoF are more likely to have uORFs (Chi-square P<1×10^−15^) and start-stops (Chi-square P=8.5×10^−05^) than genes most tolerant to LoF. The average numbers of each uAUG type across all 5’UTRs are shown by dotted lines. uORF: upstream open reading frame; oORF; overlapping open reading frame. **E**) Genes most intolerant to LoF were significantly more likely to have multiple associated CAGE peaks when compared to genes most tolerant to LoF (Supplementary Figure 5; CAGE peak >1, 91.9% vs 72.4%, Chi-square P<1×10^−15^; CAGE peak ≥6, 44.6% vs 16.3%, Chi-square P<1×10^−15^). **F**) Whilst Ribo-seq uORFs in genes intolerant to LoF appear to more frequently have canonical start-codons, this difference is not statistically significant (Chi-square P=0.18). All statistical tests compare the lowest and highest two LOEUF deciles.

Secondary structures within 5’UTRs are thought to cause inefficient ribosomal scanning [27]. The propensity of a sequence to form RNA secondary structures can be predicted from high GC content and low minimum free energy (MFE) [2,28]. We used RNAfold [29] to compute the MFE prediction per 5’UTR. The most LoF intolerant genes had lower MFE (Figure 2C: mean MFE=-115 vs -55, Wilcoxon *P*<1×10^−15^) and a higher GC content (Supplementary Figure 3A; mean=67.3% vs 59.9%; Wilcoxon *P*<1×10^−15^) than LoF tolerant genes, indicating a higher likelihood for these 5’UTRs to be structured. To demonstrate that this greater propensity to create secondary structures is over and above what would be expected given the increased length of LoF intolerant 5’UTRs (given that longer sequences have a greater propensity to create secondary structures), we repeated the analysis only on 5’UTRs between 100-300 bp in length. The results for both MFE and GC content remained significant (both Wilcoxon *P*<1×10^−15^). These results suggest that genes that are intolerant to LoF are more likely to have stable secondary structures within their 5’UTRs.

**Figure 3:**
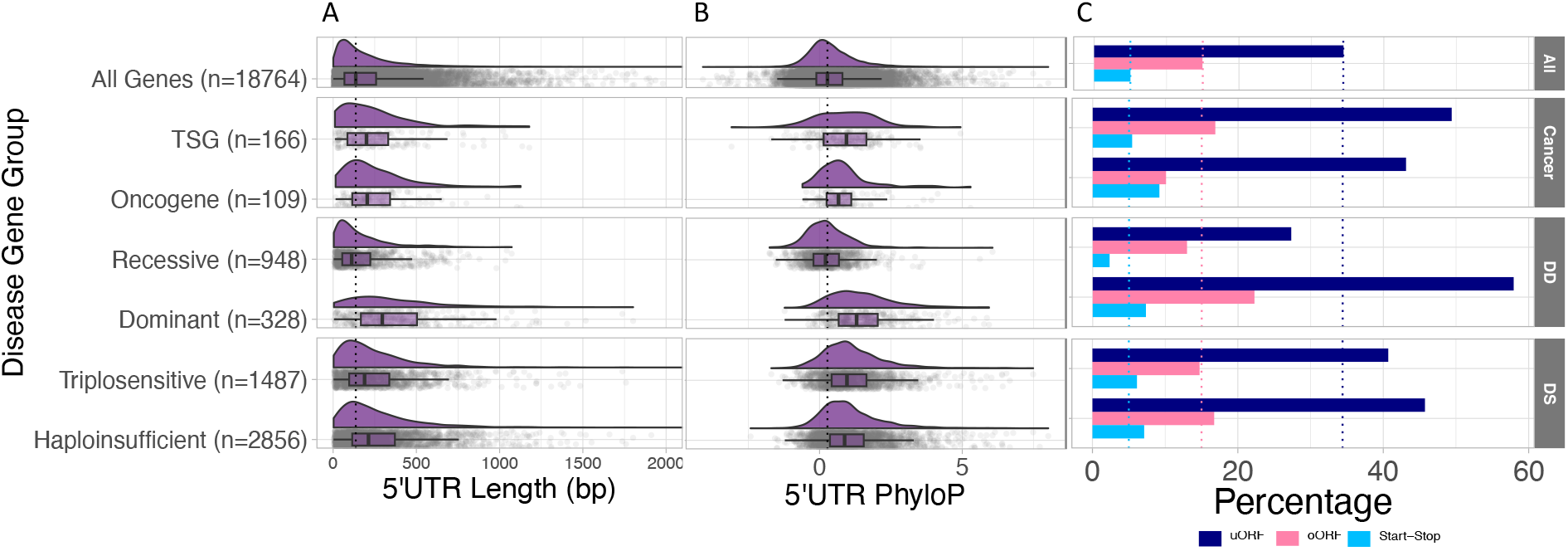
Comparison of 5’UTRs across disease genes sets. **A)** The 5’UTRs of disease genes are significantly longer (Wilcoxon: DD dominant: P<1×10^−15^; Onc: P=1.5×10^−05^; TSG:P=2.9×10^−04^; HS: P<1×10^−15^; TS: P<1×10^−15^) with the exception of DD recessive genes which are significantly shorter (Wilcoxon P=2.7×10^−08^), when compared to the average across all genes. The median 5’UTR length for all genes (136 bp) is shown by the dotted black line. The x-axis was truncated at 2,000 bp (22 genes had 5’UTRs >2,000 bp). **B)** Disease gene 5’UTRs are significantly more conserved (T-test: DD dominant: P<1×10^−15^; Onc: P=9×10^−06^; TSG: P=4.3×10^−08^; HS: P<1×10^−15^; TS: P<1×10^−15^) except DD recessive genes which are significantly less conserved (T-test: P=4.9×10^−08^), compared to all genes. The dotted black line is the median PhyloP score for all genes (0.28). **C)** Disease genes significantly more often contain uORFs (Chi-square: DD dominant: 57.9%, P<1×10^−15^; TSG=49.4%, P=6.5×10^−05^; HS=45.7%, P<1×10^−15^; TS=40.7%, P=1.4×10^−07^), when compared to all 5’UTRs. Start-stops are only significantly enriched in HS genes (P=3.1×10^−08^). The dotted lines mark the percentage of all genes with each uAUG type.

The 5’UTRs of LoF intolerant genes are more highly conserved than LoF tolerant genes, shown by significantly higher PhyloP scores [20] (Figure 2B; T-test *P*<1×10^−15^). This is even more pronounced when looking specifically at start and stop codons of predicted uORFs and start-stop elements (Figure 2B; T-test all *P*<1×10^−15^). We saw a similar pattern with Combined Annotation-Dependant Depletion (CADD) scores [21] of variant deleteriousness for all possible single nucleotide substitutions at each position, with CADD scores increasing with decreased LoF tolerance (Supplementary Figure 3C).

We next assessed the proportion of genes in each LOEUF decile with different categories of uAUGs. Genes most intolerant to LoF more frequently contain uORFs and start-stops than LoF tolerant genes (Figure 2D; 46.2% vs 27.8%; *P*<1×10^−15^, and 6.8% vs 4.5%; *P*=8.5×10^−05^ for uORFs and start-stops respectively). This is true both using predicted uORFs and the uORFs detected by Ribo-Seq (Supplementary Figure 4A). However, we would expect there to be more uAUGs in these genes as they have longer 5’UTRs. To account for this difference in 5’UTR length across deciles, we computed the number of uAUGs per base pair (bp). The 5’UTRs of the most LoF intolerant genes have significantly fewer uAUGs per bp compared to the most tolerant genes (Supplementary Figure 3D; mean=0.009 uAUG per bp vs 0.013 uAUG per bp; Chisquare *P*<1×10^−15^), suggesting that uAUGs are selectively depleted from these genes. Despite this overall depletion, 52.1% of LoF intolerant genes (bottom quintile of LOEUF) contain at least one uAUG, suggesting that they may play an important role in translational regulation of these genes.

**Figure 4:**
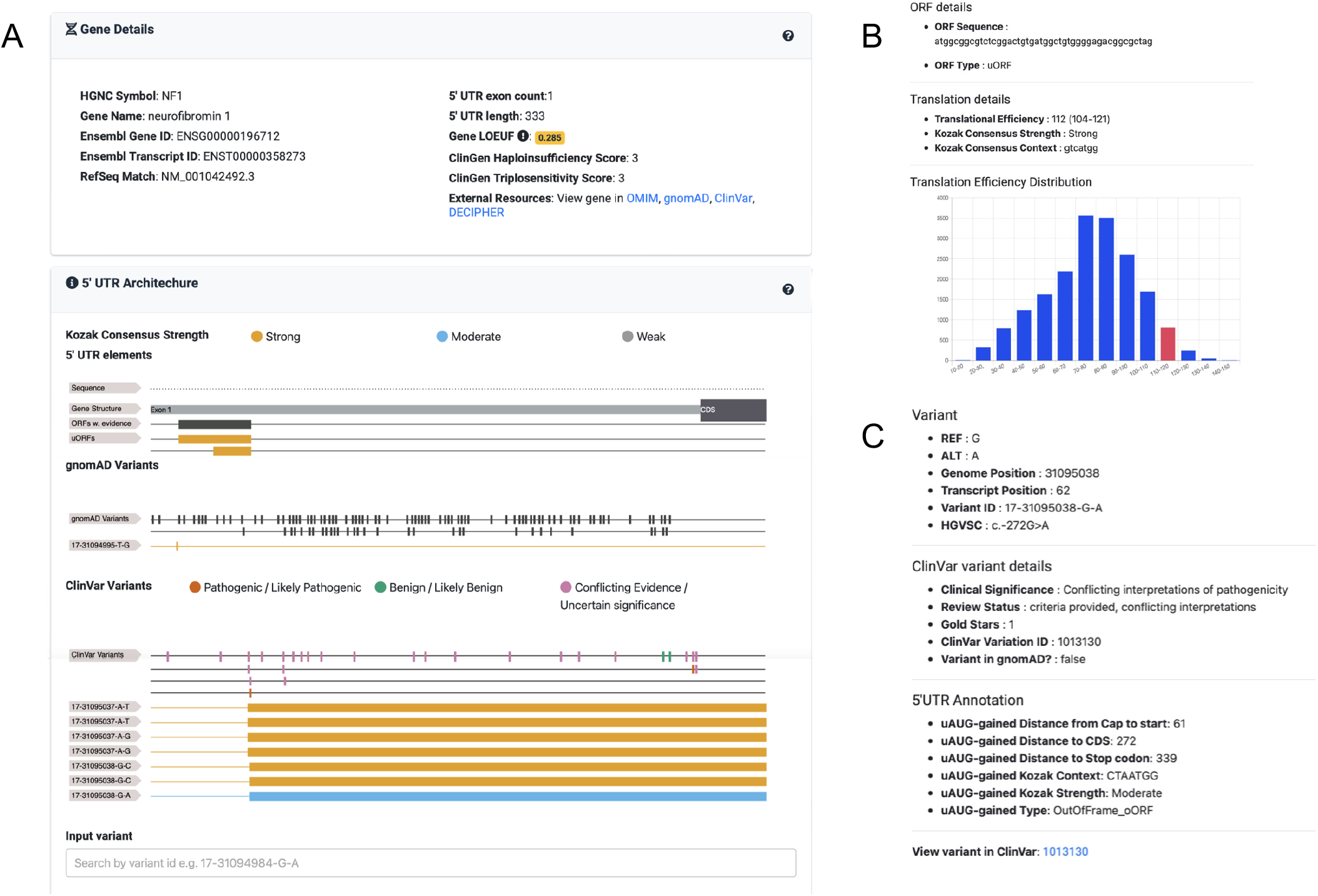
VuTR: an interactive web-based tool. **A)** A screenshot from VuTR showing the NF1 gene. The top section displays summary gene details and links to other tools and databases. The following section, titled ‘5’UTR Architecture’, shows the gene’s native 5’UTR exon structure, predicted uAUG elements, and Ribo-Seq uORFs. NF1 features two predicted uORFs, both with a strong Kozak consensus strength (shown by the yellow colour), the longer 45 bp uORF is also found in the Ribo-Seq dataset. Variants observed in gnomAD and ClinVar are displayed in separate tracks. Each variant that creates a uORF or disrupts a predicted uORF is shown on a separate row. Here, a variant that disrupts the start of the longer predicted uORF, which is also found in the Ribo-Seq data (uAUG-lost; 17-31094995-T-G) is observed in gnomAD. Four ClinVar variants create out-of-frame ORFs (oORFs) by either disrupting the stop codon of the two native uORFs (uSTOP-lost; 17-31095037-A-T, 17-31095037-A-G, 17-31095038-G-C) or by creating a new oORF through a uAUG-gained variant (17-31095038-G-A). **B)** An example of a popup that appears when a uORF / oORF is selected, giving context specific details regarding its sequence, Kozak consensus strength, and a histogram of how its predicted translational efficiency [6] compares to all other uORFs / oORFs within MANE 5’ UTR sequences. **C)** An example of a popup that appears when a ClinVar variant is selected. This example shows the uAUG-creating variant, 17-31095038-G-A.. The popup displays the variant details, information from ClinVar, and the variant annotation from UTRannotator [35]. uORF: upstream open reading frame.

To determine whether the likelihood of uORF translation, and hence strength of repression of downstream CDS translation, differed between LOEUF deciles we compared the start contexts of predicted uORFs to a dataset of experimentally measured translational efficiencies (TE), quantified across a range of cell lines [6]. We saw no significant difference in TE of uAUGs across deciles (Supplementary Figure 4B, Wilcoxon *P*=0.6), nor a significant enrichment of canonical over non-canonical start site usage of the Ribo-Seq uORFs (Figure 2F; Chi-square, *P*=0.18).

Whilst we have used the MANE Select transcript set to limit our above analysis to a single, representative transcript per gene, alternative transcription start site (TSS) usage is a major contributor to transcript isoform diversity and gene regulation [30]. Cap Analysis of Gene Expression (CAGE) tags the 5’ ends of mRNA transcripts, allowing us to analyse alternative TSS usage. To observe the diversity of 5’UTRs across the LOEUF spectrum, we used CAGE data from the FANTOM consortium [31]. Genes most intolerant to LoF were significantly more likely to have multiple associated CAGE peaks when compared to genes most tolerant to LoF (Figure 2E; CAGE peak >1, 91.9% vs 72.4%, Chi-square *P*<1×10^−15^; CAGE peak ≥6, 44.6% vs 16.3%, Chi-square *P*<1×10^−15^). Assessing alternative splicing possibilities, we found no significant difference in the proportion of genes that have 5’UTR introns across LEOUF deciles (Supplementary Figure 3B, Chi-square *P*=0.19).

Finally, we hypothesised that the uORFs in LoF intolerant genes might be optimised to promote efficient uORF translation and re-initiation at the CDS start-codon. We assessed codon optimality (tAI scores) of the Ribo-Seq uORFs, but found no significant differences between deciles (Supplementary Figure 5A; Wilcoxon *P*=0.17). We observed a very small, but significant difference in average uORF length across deciles (means 52.5 bp vs 59.1 bp, Wilcoxon *P*=4.9×10^−06^), but only when considering the predicted uORF and not the Ribo-Seq set (Supplementary Figure 5B, 5C; Wilcoxon *P*=0.9). We also observed that the stop codons of the uORFs closest to the CDS start are significantly further upstream of the CDS start in more LoF intolerant genes (Supplementary Figure 5D; means 99 bp vs 77 bp, Wilcoxon *P*=1.3×10^−04^). In other words, these genes have a greater potential re-initiation distance.

### Translational regulation through 5’UTRs is important for genes involved in disease

Given the increased complexity of 5’UTRs observed in LoF intolerant genes, we were interested to see whether these results were relevant to 5’UTRs of genes where disruption of tight regulatory control may lead to disease. We investigated 5’UTR features in genes within which predicted LoF variants have been reported to cause developmental disorders (DD) and cancer, as well as a wider set of dosage sensitive (DS) genes [32–34]. For DD genes, we compared dominant and recessive genes, given the former are more likely to be highly dosage sensitive. For cancer, we analysed tumour suppressor genes (TSGs) and oncogenes separately (Onc). Finally, for DS genes we compared haploinsufficient (HS) and triplosensitive (TS) genes. For all statistical tests we compared the disease gene group against all MANE Select 5’UTRs with that specific disease group removed.

Whilst 5’UTRs average 202 bp in length, disease gene 5’UTRs are significantly longer (Figure 3A; DD dominant: 369 bp, Wilcoxon *P*<1×10^−15^; Onc: 260 bp, Wilcoxon *P*=1.5×10^−05^; TSG: 254 bp, Wilcoxon *P*=2.9×10^−04^; HS: 279 bp, Wilcoxon *P*<1×10^−15^; TS: 253 bp, Wilcoxon *P*<1×10^−15^). A significantly higher number of disease gene 5’UTRs contain uORFs than the average of 34.4% across all genes (Figure 3C; DD dominant: 57.9%, Chi-square *P*<1×10^−15^; TSG=49.4%, Chi-square *P*=6.5×10^−05^; HS=45.7%, Chisquare *P*<1×10^−15^; TS=40.7%, Chi-square *P*=1.4×10^−07^), although the difference is not-significant for the oncogene gene set (43.1%, Chi-square *P*=0.07). Start-stop elements are only significantly enriched in HS genes (7.1% vs 5.0%; Chi-square *P*=3.1×10^−08^), however, given the small number of genes that contain start-stops, we are likely underpowered to detect a significant enrichment in our smaller gene sets.

Disease gene 5’UTRs are also significantly more conserved when compared to all genes (Figure 3B; DD dominant: T-test *P*<1×10^−15^; Onc: T-test *P*=9×10^−06^; TSG: T-test *P*=4.3×10^−08^; HS: T-test *P*<1×10^−15^; TS: T-test *P*<1×10^−15^). We did not observe a significant difference in the number of 5’UTR introns between disease gene sets and all genes (Supplementary Figure 6; DD dominant: Chi-square *P*=0.07; Onc: Chi-square *P*=0.09, TSG: Chi-square *P*=0.15; HS: Chi-square *P*=0.18; TS Chi-square *P*=0.39).

We observed a marked distinction between DD dominant and recessive gene 5’UTRs. When compared to the average across all genes, the 5’UTRs of DD recessive genes were significantly shorter (Figure 3A; mean=169 bp, Wilcoxon *P*=2.7×10^−08^), have significantly fewer 5’UTR introns (Supplementary Figure 5B; Chi-square *P*=4.7×10^−06^), are significantly less conserved (Figure 3B; PhyloP, T-test *P*=4.9×10^−08^), and also have fewer uORFs and start-stops (Figure 3C; Chi-square *P*=2.7×10^−06^ (uORFs); Chi-square *P*=1.3×10^−04^ (start-stops)). The lower complexity of the 5’UTRs of this recessive gene set likely reflects their insensitivity to changes in dosage. The observation that these 5’UTRs are significantly different to the all gene average likely reflects the fact that the all gene set contains many genes that are sensitive to dosage changes. To account for this, we tested DD recessive genes against genes in the middle two LEOUF deciles; we see no significant difference in 5’UTR length (mean 169 vs 177 bp, Wilcoxon *P*=0.04), the number of uORFs (Chi-square *P*=0.66), or mean PhyloP scores (T-test *P*=0.1). We do still observe significantly fewer introns in DD recessive genes (Chi-square *P*=8.2×10^−05^).

### Visualising 5’UTRs with VuTR

Here, we have presented an overview of 5’UTRs across different gene sets, however, there is still considerable variability within each set. To support investigation of individual gene 5’UTRs, their regulatory features, and genetic variation within them, we have created an interactive web-based tool, VuTR (pronounced view TR; https://vutr.rarediseasegenomics.org/). For a query gene symbol or MANE transcript ID, VuTR displays the sequence of the 5’UTR, statistics including the length and number of uAUGs, and the distribution of both predicted and Ribo-Seq uORFs within the 5’UTR. Further, VuTR uses annotations from UTRannotator [35] to display variants in gnomAD [24] and ClinVar [36] that create uAUGs or disrupt predicted uORFs. Figure 4 shows the output for *NF1*.

## Discussion

Here, we characterised the features of 5’UTRs across all human genes to understand the natural variability in these regions. We further investigated the differences in 5’UTR composition across deciles of tolerance to LoF and between sets of disease genes. Our findings show that genes sensitive to LoF have significantly different 5’UTRs; they are longer, more conserved, have higher propensity to be structured, and contain more uORFs, than genes that are tolerant to LoF.

The increase in length and complexity of the 5’UTRs of dosage sensitive genes points to the importance of post-transcriptional/translational regulation in controlling the levels of encoded proteins. This is further supported by the stark difference we observed between DD dominant and DD recessive genes, where recessive genes that are not sensitive to changes in dosage have shorter 5’UTRs with less complexity. We observe increased length and complexity across both haploinsufficient and triplosensitive gene sets, although we acknowledge that there is considerable overlap between these sets.

This work aimed to provide a general picture of the variation in 5’UTR complexity, but it has several limitations. We only analysed a single transcript per gene; we used the highly curated MANE Select transcript set, which likely reflects the most clinically relevant transcript per gene. We acknowledge there are other relevant transcripts that we have not included. To mitigate not accounting for complexity at the level of alternative 5’UTR isoforms we used CAGE data to determine the number of TSS’s per gene, however, this only assess differences in TSS usage and not alternative splicing within 5’UTRs derived from the same TSS.

We used two different uORF sets throughout this work, a predicted set derived from every AUG within each 5’UTR, and an experimental set from Ribo-Seq [17]. Our predicted uORF set likely contains many uORFs that are not translated. Conversely, due to necessary stringent filtering, and tissue and temporal specificity of uORFs, there are likely many uORFs that are translated, but that are not captured in the Ribo-seq data we included. Other work has also shown preferential uORF usage under stress conditions [37,38]. Our predicted uORF set is also only based on canonical start sites, whereas 45.3% of the Ribo-seq uORFs use non-canonical start sites. Therefore, there are likely many more potentially translated uORFs which are excluded from our predicted uORF set. Despite these limitations, our results are consistent across both the predicted and experimental uORF sets.

Here we have focussed on uORFs as *cis* regulators of translation, however, there is evidence from mass spectrometry that some uORFs encode a detectable peptide product (SEPs; smORF encoded peptides) [39]. Other work has demonstrated that some SEPs may have a biological function [40]. Further work needs to be done to find and curate these and to understand their role.

We limited this work to analysis of 5’UTRs, however, these are only a fraction of the overall mRNA transcript. The wider mRNA length and composition plays an important role in transcript stability and secondary structure. Further work is needed to jointly analyse 5’UTR and 3’UTR elements.

We have analysed broad trends in 5’UTRs across gene categories, but there remains considerable variety within each category. For example, whilst the 5’UTRs of LoF intolerant genes tend to be much longer than average, some LoF intolerant and known dosage sensitive disease genes have very short 5’UTRs. For example the 5’UTR of *FOXF1*, a haploinsufficient DD gene which is in the 2nd LEOUF decile, is only 43 bp long. LoF variants in *FOXF1* are a known cause of alveolar capillary dysplasia with misalignment of pulmonary veins. This variability may limit attempts to use the 5’UTR features to predict gene dosage sensitivity and points to a much more complex regulatory landscape. We have created the open-source web-tool VuTR to enable investigation of 5’UTRs of specific genes.

Here, we have assessed how 5’UTRs vary by gene tolerance to LoF. Overall, our work supports the important role of 5’UTRs in tightly regulating protein levels, particularly in genes that are sensitive to changes in dosage. This increased knowledge of 5’UTR diversity will aid interpretation of genetic variants in 5’UTRs for a role in disease.

## Methods

### Defining and annotating a high-confidence set of 5’UTRs

We used MANE Select transcripts from v1.0 of the MANE resource [26] to define a single 5’UTR per gene. Of 19,062 MANE Select transcripts, 18,764 had annotated 5’UTRs.

5’UTR length was calculated as the total length of all exons for each 5’UTR.

The GC content of each 5’UTR was calculated by dividing the number of G and C bases by the length of the 5’UTR.

5’UTR bases were further annotated with per-base vertebrate PhyloP scores (phyloP100way) retrieved in R using GenomicScores package and Combined Annotation Dependant Depletion (CADD) v1.6 using the CADD version 2.2.0 release files and tabix (HTSlib v1.9: foss/2018b) to filter MANE 5’UTR coordinates.

### Identifying and classifying uAUGs

We identified all ATGs in the sequence of each 5’UTR as upstream AUGs (uAUGs). Each uAUG was then annotated as one of the following categories:

1. As a start-stop, if the uAUG was immediately followed by a stop codon.
2. As a uORF if there was an in-frame stop codon (TAA, TAG, TGA) within the 5’UTR.
3. As an oORF if there was no in-frame stop codon within the 5’UTR. These were further subdivided into out-of-frame oORFs if the uAUG was not in-frame with the CDS, or in-frame n-terminal extensions (NTEs) if the uAUG was in-frame to the CDS.

Translational efficiencies (TE) of uAUGs were determined using work by Noderer *et al*., 2014 [6] by matching to the surrounding sequence context. They used fluorescence-activated cell sorting and high-throughput DNA sequencing (FACS-seq) to determine efficiency of start codon recognition for all possible translation initiation sites using AUG start codons, across a variety of cell lines.

Where the uAUG TE sequence was not complete as too close to the start of the 5’UTR, these uAUGs were excluded from this analysis.

### Defining a set of uORFs with experimental evidence

Ribo-seq data from Chothani et al. [17] was downloaded from https://smorfs.ddnetbio.com/ and filtered to include only uORFs.

To determine the codon optimality of Ribo-Seq uORFs, we used previous work based on tAI (tRNA adaptive indices) in HeLa cells [42]. This scores each codon as “optimal” or “not-optimal”. Each codon in a Ribo-seq uORF was translated into whether it was optimal (noted as 1) or not (noted as 0). Adding these numeric codons, we then divided by the total number of codons for each uORF to get a total optimality score; with higher scores being more optimal.

### Categorising 5’UTRs into deciles of LoF tolerance

LOEUF scores were downloaded from gnomAD (v2.1.1). We filtered to the canonical transcript and where genes had multiple LEOUF scores we kept the transcript with the higher score. They were then binned into deciles. We then matched each gene to the MANE set based on Ensembl stable gene id’s.

### Identifying disease-gene sets

Developmental disorder genes were downloaded (18 February 2021) from DDG2P [32]. They were limited to confirmed or probable roles in developmental disorders and loss-of-function disease mechanism.

The COSMIC Cancer Gene Census [33] was downloaded 22nd February 2021. They were restricted to nonsense, frameshift and missense mutation types and then filtered to oncogene or TSG only as cancer gene type.

Dosage sensitive genes (haploinsufficient and triplosensitive) gene sets were taken from the work by Collins *et al*. [34].

### Calculating minimum free folding energies of 5’UTRs

The Vienna RNA package was downloaded 28 July 2022 (https://www.tbi.univie.ac.at/RNA/index.html) and used the RNAFold v2.5.1 program on 5’UTR full exon sequences to predict the minimum free energy secondary structure.

### Assessing transcription start site (TSS) diversity

Data downloaded from FANTOM5 “CAGE peak based annotation table of robust CAGE peaks for human samples” (30 November 2022). We used CAGE peaks which uniquely associate to a gene. CAGE data only included HGNC id’s so these were used to match with MANE genes.

### Creating an interactive web-based 5’UTR visualisation tool

VuTR’s front end uses the AdminLTE (https://adminlte.io/) template. Its main gene page utilises the FeatureViewer (http://calipho-sib.github.io) to visually display tracks for genes, variants and any native, or altered ORFs. ChartJS (http://ChartJS.org/) is used for plotting web charts. The backend of VuTR was built using Flask as a web framework and Flask-SQLAlchemy as an object-relational mapping tool to connect with SQLite3 databases. The application was wrapped within a Docker python:3.9.7-slim-buster base image and served using nginx/1.18.0 reverse-proxy on Ubuntu 22.04.1. VuTR is available at http://VuTR.rarediseasegenomics.org/ and is released under the GPL version 2 licence. The code is available at https://github.com/Computational-Rare-Disease-Genomics-WHG/VuTR where a list of additional packages can be found.

VuTR uses MANE v1.0 transcripts. Genes were matched to LOEUF scores and with ClinGen Haplo- and Triplosensitive data from https://ftp.clinicalgenome.org/ClinGen_gene_curation_list_GRCh38.tsv. Predicted ORFs were annotated with their Kozak consensus sequences, lengths and locations. We then matched each ORF with its translational efficiency dataset from Noderer et al., 2014 [6]. All datasets were linked using their stable Ensembl gene identifiers where available and then ingested into an SQLite3 database.

Additionally, a separate variant-specific SQLite database was produced. Here using the MANE v1.0 cDNA sequences, a set of all possible single nucleotide variants, and small indels (up to 3 bp in length) were generated within 5’ UTR exons. We then annotated these variants with their variant effect using the Ensembl Variant Effect Predictor Version 103 with the UTR annotator plugin [35,43]. Additionally, this set was flagged if any variants also appeared in gnomAD v3.1.1 and within ClinVar Weekly release.

### Statistical tests

To account for multiple testing we calculated a study-wide *P*-value threshold of 8.3×10^−4^ using a Bonferroni correction based on 60 statistical tests. All *P*-values less than 1×10^−15^ are reported as *P*<1×10^−15^.

## Declarations

### Ethics approval and consent to participate

Not applicable

### Consent for publication

Not applicable

### Availability of data and materials

The datasets analysed are available in the Computational Rare Disease Genomics Github (https://github.com/Computational-Rare-Disease-Genomics-WHG/5-UTR_characterisation)

### Authors’ contributions

Analyses were led by NWieder with contributions from END, ACM-G, FHL, JT-M and MF. SPC, OJLR and SS contributed data. JLA, DGM, and RWD critically evaluated the work and provided feedback. The project was conceived and supervised by NWhiffin. All authors read and approved the final manuscript.

## Acknowledgements

None

