## Supplementary material for "Differences in 5’untranslated regions highlight the importance of translational regulation of dosage sensitive genes": supp figures

### Supplementary Figures

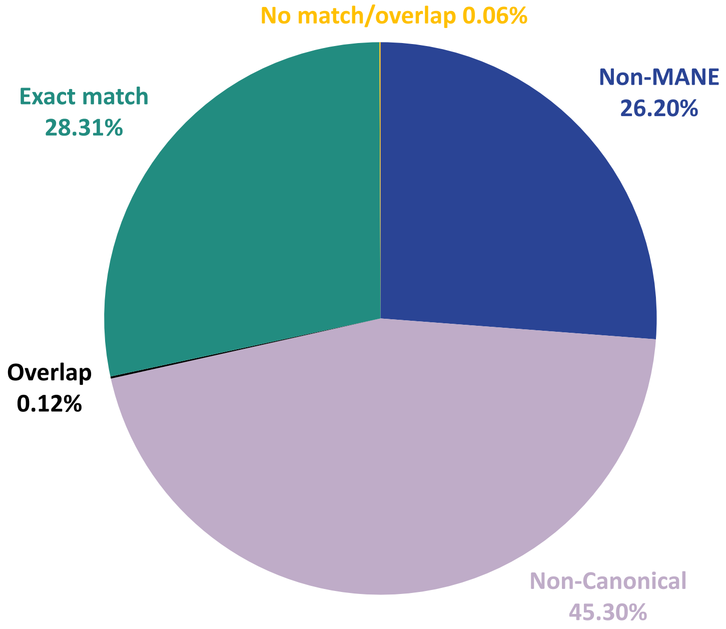


**Supplementary Figure 1**: *Overlap between Ribo-seq uORFs (n=5,052) and predicted MANE uORFs (n=18,064). 28.3% of Ribo-seq uORFs matched a predicted uORF exactly. A further 0.12% overlapped with a predicted uORF. 26.2% of Ribo-seq uORFs did not map to MANE transcript 5'UTRs so we did not include in our analyses.45.3% of Ribo-seq uORFs start with a non-canonical start codon (non-AUG) and hence would not be predicted to overlap with the predicted uORF set.*

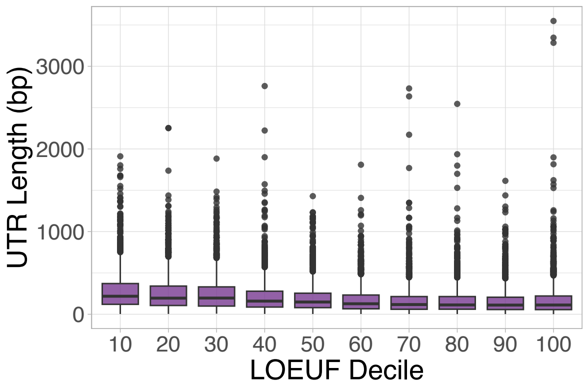


**Supplementary Figure 2**: *5'UTR length by LOEUF decile with genes within bottom 10% of coding sequence length distribution removed.*

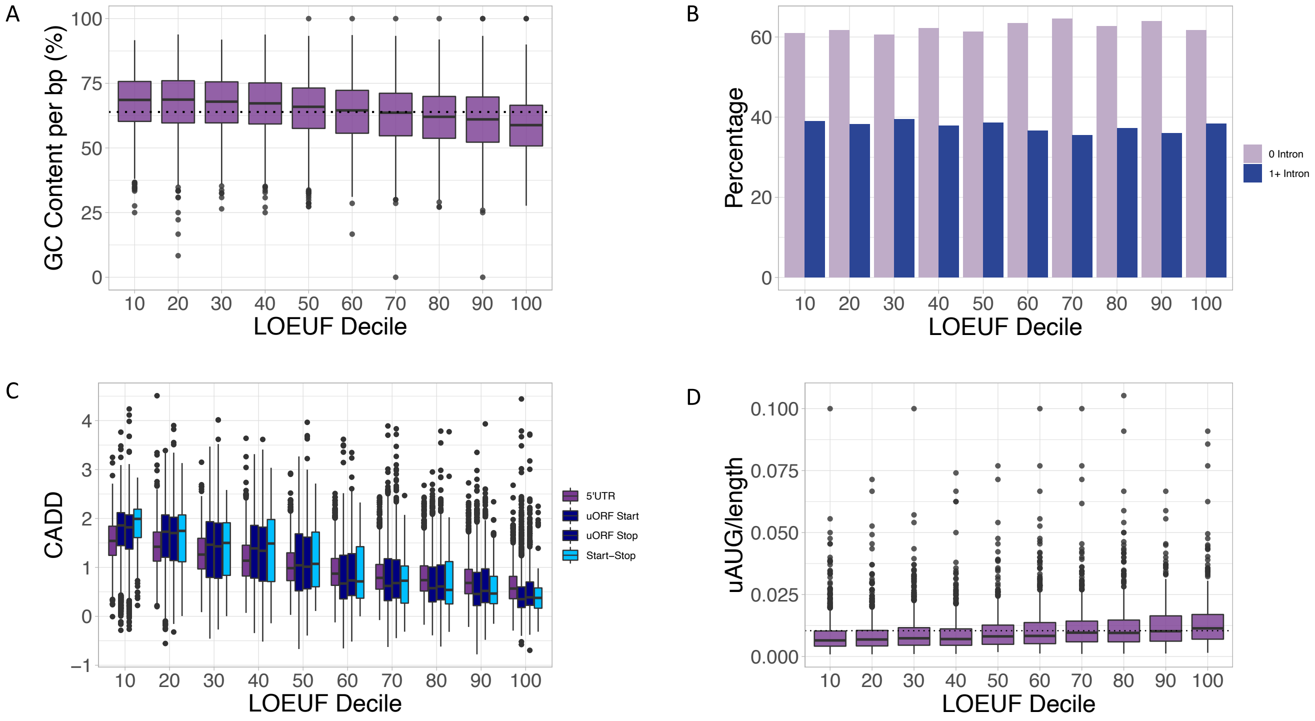


**Supplementary Figure 3**: ***A****) Genes most intolerant to LoF had higher GC content than LoF tolerant genes (Wilcoxon $P<1x10^{-15}$). Average GC content for all genes is 63.9% and is shown by a dotted line.* ***B****) There was no significant difference between 5'UTR intron proportions across LEOUF deciles (Chi-square P=0.19).* ***C****) CADD scores for 5'UTRs, uORFs and start-stops across LEOUF deciles were higher in lower deciles; in genes more intolerant to LoF (T-test 5'UTRs $P<1x10^{-15}$, uORF start $P<1x10^{-15}$, uORF stop $P<1x10^{-15}$, start-stop $P<1x10^{-15}$).* ***D****) Genes most intolerant to LoF had fewer uAUGs per base pair (Chi-squareP$<1x10^{-15}$). Average uAUG/length is 0.01 (dotted line). All statistical tests compare the bottom two and top two deciles of LOEUF.*

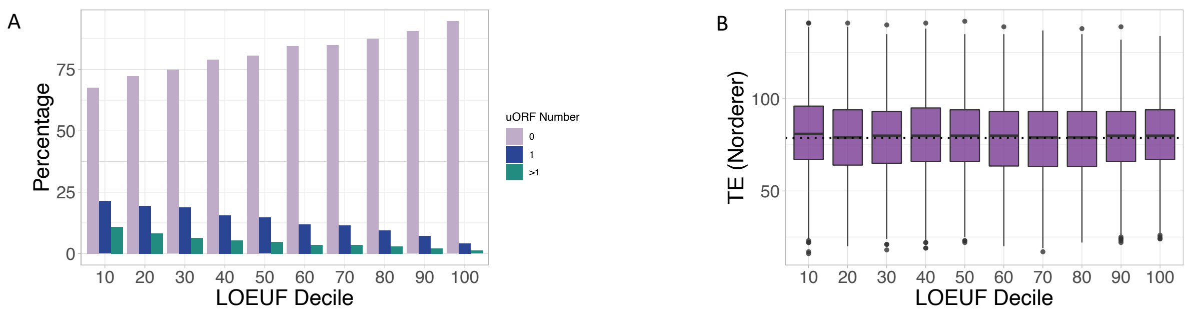


**Supplementary Figure 4**: ***A****) The 5'UTRs of genes that are more intolerant to LoF more often contain Ribo-seq uORFs than genes that are tolerant to LoF (Chi-square P<1x10$^{-15}$).* ***B****) There was no difference between translational efficiencies (TE) for predicted uAUGs across LEOUF deciles (Wilcoxon P=0.6). Statistical tests compare the bottom two and top two deciles of LOEUF.*

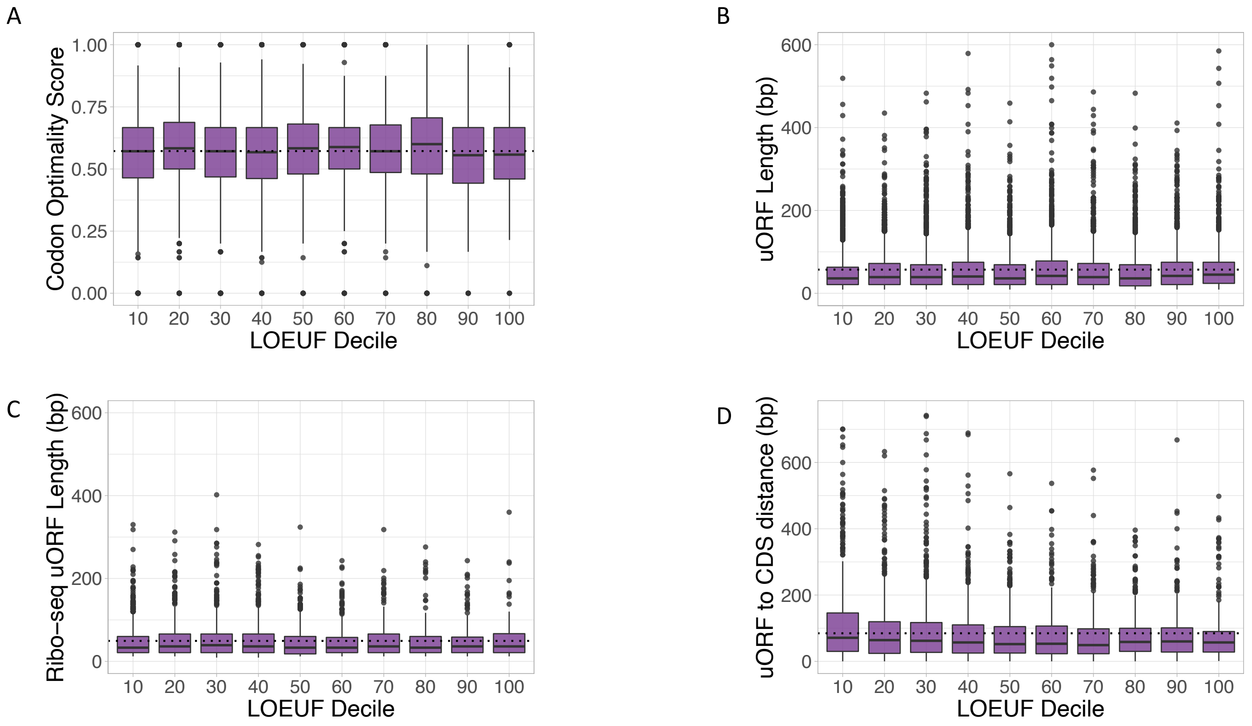


**Supplementary Figure 5**: *A) There was no significant difference in codon optimality score for Ribo-seq uORFs between lowest and highest two deciles (Wilcoxon P=0.17), average for all genes is dotted line.* ***B****) Predicted uORFs in the lower two deciles were slightly shorter compared to the highest two deciles (Wilcoxon $P=4.9x10^{-06}$). Mean uORF length is shown as a dotted line. The y-axis is truncated at 600bp (7 uORFs were >600bps).* ***C****) There is no significant difference in Ribo-seq uORF lengths between lowest and highest two LOEUF deciles(Wilcoxon, P=0.9). Mean uORF length is shown as a dotted line. The y-axis is truncated at 600bps (1 uORF was >600bps).* ***D****) The distance between the stop codon of the last uORF to the CDS start (of the predicted uORF set) was greater in the genes most intolerant to LoF (lowest versus top two deciles, Wilcoxon $P=1.3x10^{-04}$). Average distance is 84.7bps (dotted line). The y-axis is truncated at 750bp (3 genes had distance >750bps).*

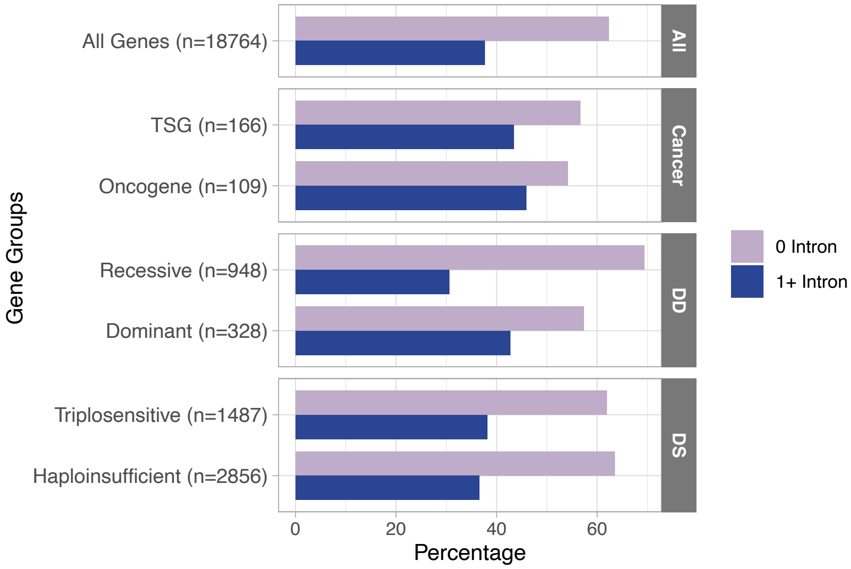


**Supplementary Figure 6**: *We did not observe any difference in the proportion of 5UTRs that have introns between disease gene sets and all genes (DD dominant: Chi-square P=0.07; Onc Chi-square P=0.09; TSG Chi-square P=0.15; HS Chi-square P=0.18; TS Chi-square P=0.39), except for DD recessive genes, where significantly fewer had introns (Chi-square P=4.7x10$^{-06}$).*
